# Trans-NanoSim characterizes and simulates nanopore RNA-seq data

**DOI:** 10.1101/800110

**Authors:** Saber Hafezqorani, Chen Yang, Ka Ming Nip, René L Warren, Inanc Birol

## Abstract

We introduce Trans-NanoSim, the first tool that simulates reads with technical and transcriptome-specific features learnt from nanopore RNA-seq data. Through benchmarking on sets of nanopore reads from human and mouse reference transcriptomes, we show the robustness of Trans-NanoSim in capturing the characteristics of nanopore cDNA and direct RNA reads. As a cost-effective alternative to sequencing real transcriptomes, Trans-NanoSim would facilitate the rapid development of analytical tools for nanopore RNA-seq data. Trans-NanoSim is freely accessible at https://github.com/bcgsc/NanoSim

## Background

RNA-sequencing (RNA-seq) is a cornerstone technology that has helped study and further our understanding of transcriptomes [1]. Third-generation single-molecule sequencing technologies such as those from Oxford Nanopore Technologies (ONT, Oxford, UK) are proving invaluable for isoform-level analyses. For example, ONT reads, ranging from 1-100 kb long, permit identification and quantification of most full-length isoforms in the human transcriptome and enable various complex feature analyses [2–5]. In recent years, there has been an increase in development of novel algorithms to leverage the power of this technology [6–11]. In this active field, simulated data with a known ground-truth is a cost-effective means to help develop, refine, and benchmark these tools.

Long-read simulators have been developed for ONT genomic reads [12–13]. For example, NanoSim [12] utilizes statistical models to learn the characteristics of user-provided data, and then applies those models to simulate ONT genomic reads. DeepSimulator [13] employs a context-dependent deep learning model to simulate the electrical current signal, followed by base-calling. However, neither tool is specifically designed to simulate transcriptomic reads. Further, they do not address transcriptome-specific features such as transcript expression profiles and intron retention (IR) events. While transcript expression levels inform about the biological state of a transcriptome, IR, as one of the main forms of alternative splicing, contributes to the functional complexity of eukaryotic transcriptomes [14]. ONT reads have the potential in capturing complex IR events involving multiple introns, thus allowing researchers to investigate IR at isoform-level resolution. Currently, there is an unmet need for an ONT RNA-seq simulator, which can aid the development of transcriptome analysis methods without the expense of sequencing experiments.

Here we present Trans-NanoSim, the first ONT transcriptome read simulator, implemented within the NanoSim [12] package. When provided with experimental data, Trans-NanoSim first learns platform-specific features including read length distributions and error models, and transcriptome-specific features including IR events and expression profiles. Then, at the simulation stage, it utilizes these profiles to generate *in silico* reads for a given reference transcriptome (**Fig 1**, **Methods**). We benchmarked the performance of Trans-NanoSim against DeepSimulator by generating sets of synthetic reads using publicly available experimental ONT cDNA and direct RNA reads describing human and mouse transcriptomes (**Methods**, **Supplementary Note 1** and **2**).

**Fig 1:**
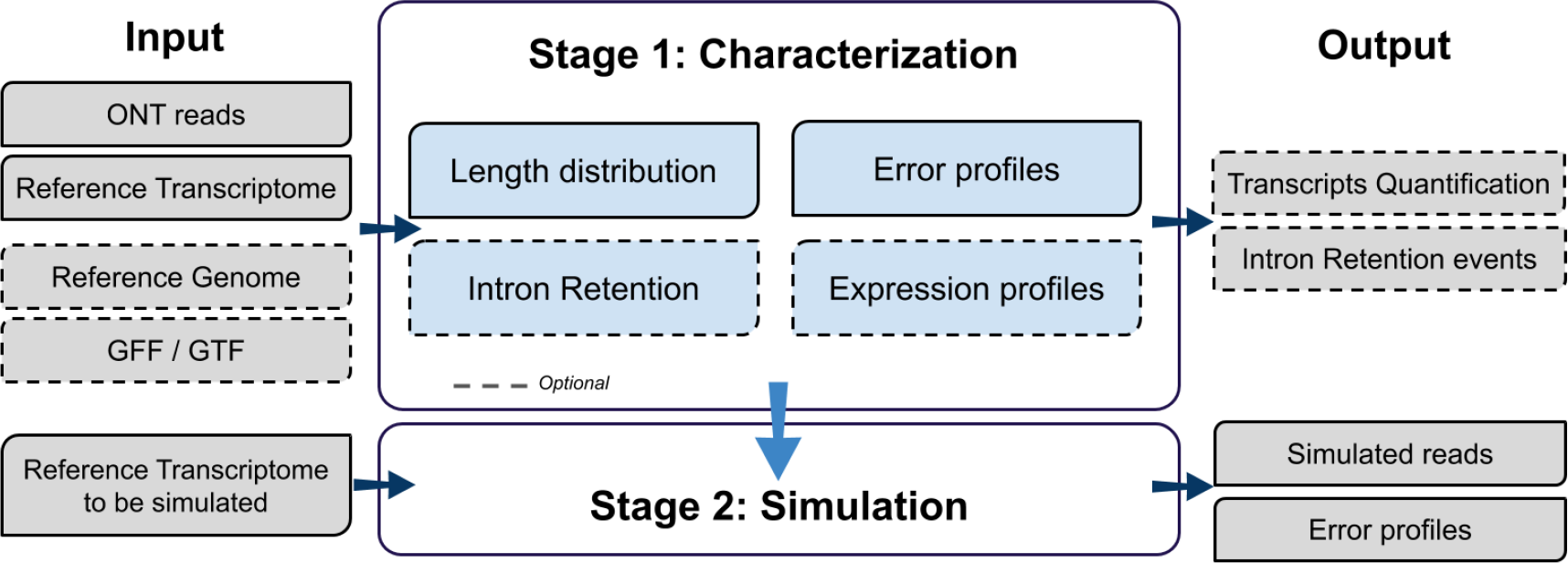
Schematic overview of the Trans-NanoSim pipeline. The first stage (Characterization) of the pipeline aligns input ONT transcriptome reads against the reference transcriptome to statistically model the read length distribution and error modes. It also optionally detects intron retention events, and quantifies transcript expression. The length, error and expression profiles are then used in the second stage (Simulation) to generate synthetic reads, also reporting their associated error profiles.

## Results and discussion

For the human cDNA dataset, the length distribution of synthetic reads generated by Trans-NanoSim (mean = 853 nt) followed the empirical read length distribution (mean = 811 nt) closely (**Fig. 2a**). Through bootstrapping, the mean length of synthetic reads remained consistent with a standard deviation of 1 nt (**Supplementary Note 3**). Although we configured DeepSimulator to preserve the mean read length of empirical reads, the mode of synthetic read lengths was ~150 nt. We believe that this limitation is due to the predefined read-length distributions of DeepSimulator. Further, being a genome simulator, it does not associate the isoform expression levels with the lengths of constituent reads.

**Fig 2:**
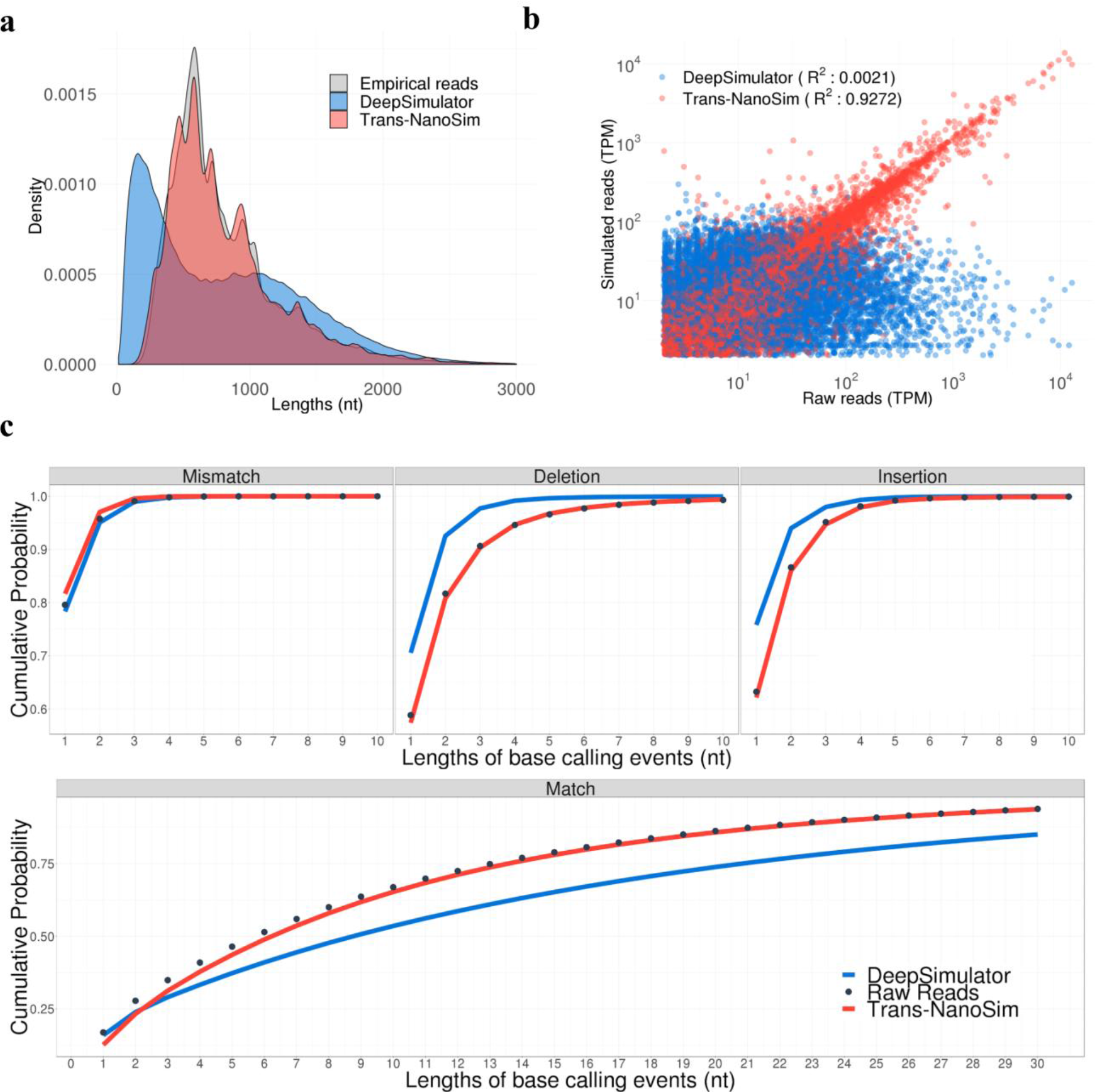
Benchmarking Trans-NanoSim and DeepSimulator on the human cDNA dataset. **a,** Comparison of length distributions of experimental reads and simulated reads generated by Trans-NanoSim and DeepSimulator. **b,** Transcript expression levels measured from simulated reads versus the same measured from experimental reads. **c,** The length of consecutive match/error bases of empirical and simulated reads, as indicated.

To determine whether synthetic reads generated by both tools account for transcript isoform expression and usage, we used the *quantify* module in Trans-NanoSim to measure the transcript expression levels for both empirical and simulated reads from a human reference transcriptome (**Methods**, **Supplementary Note 2**). The coefficient of determination (R^2^) between the estimated transcript abundance of empirical ONT reads and the simulated reads generated by Trans-NanoSim and DeepSimulator were 0.9272 and 0.0021, respectively (**Fig. 2b**). These results indicate that reads simulated by Trans-NanoSim follow the expression profiles of the empirical reads closely, which is a feature not available in genomic simulators.

The IR modeling module of Trans-NanoSim identified 2,594 transcripts in the human cDNA dataset with at least one retained intron, and nearly half of them (1,238 transcripts) were expressed at over two Transcripts Per Million (TPM) (**Methods**, **Supplementary Note 2**). Interestingly, a transcript containing five retained introns (Ensembl transcript ID: ENST00000290541.7) was highly expressed (TPM = 389). The IR modeling module also reports the transitional probability of each intron being retained based on the state of previous intron, a model that the pipeline uses for read simulations. In the human cDNA dataset, only 0.29% of reads spanned the first intron of the represented transcript. However, given an intron is retained, the probability of observing the subsequent intron being retained was 10.78%.

Finally, we aligned the synthetic and empirical reads to the reference transcriptome, and evaluated the length of consecutive match/error bases in both sets (**Methods**, **Supplementary Notes 2**). While the error rate of the empirical ONT reads from human cDNA dataset was 15.30%, the synthetic reads generated by Trans-NanoSim and DeepSimulator were 14.10% and 9.19%, respectively (**Supplementary Table 1**). Combined with the length distribution of base-calling events, it is evident that Trans-NanoSim mimics error and match events more closely to the experimental data (**Fig. 2c**).

We evaluated the computational performance of Trans-NanoSim and DeepSimulator through characterizing and simulating 890,503 reads describing the human reference transcriptome (**Supplementary Note 2**). Although both tools allow users to train their model with any dataset, authors of DeepSimulator noted that this step is computationally intensive, and advised their users against trying it (https://github.com/lykaust15/DeepSimulator). On the contrary, Trans-NanoSim took merely an hour to train. In simulation stage, Trans-NanoSim ran for 19h50m with peak memory of 990 MB while DeepSimulator ran for 45h02m with peak memory of 17.22 GB. Trans-NanoSim performed even faster (6h38m) when IR modeling is not requested (Additional information in **Supplementary Table 2**).

We recapitulated our result by repeating all analysis presented here on human direct RNA and mouse cDNA sequencing data and obtained similar findings (**Supplementary Figs. 1** and **2**, respectively). Our evaluations demonstrate the robustness of Trans-NanoSim in learning and mimicking the length distribution and sequence error profiles of nanopore RNA-seq reads. Moreover, Trans-NanoSim provides a solution to the characterization of transcriptome-specific features, such as isoform expression and IR events, which cannot be addressed by genomic read simulators. As the first ONT RNA-seq read simulator, Trans-NanoSim will offer important resources to the community. We anticipate that it will facilitate the development of novel isoform-level analysis algorithms, including transcriptome assemblers and aligners that leverage the potential of long nanopore reads.

## Acknowledgements

We thank Jared Simpson for his contribution to the transcript expression level quantification module of the pipeline.

## Authors’ contributions

SH and CY contributed equally to this work. IB, SH, and CY conceived and designed the study. SH and CY implemented the algorithm with the help of KMN and RLW. SH drafted and all the other authors reviewed, edited, and approved the final manuscript.

## Methods

### Trans-NanoSim workflow

The main workflow of Trans-NanoSim consists of two stages: (1) characterization and (2) simulation (Figure 1). In the characterization stage, Trans-NanoSim utilizes statistical models to profile read error modes, read length distribution, transcript expression patterns, and intron retention (IR). The first step in characterization stage is to align a set of input reads against a reference transcriptome, using minimap2 [7] by default (see Supplementary Note 2 for version numbers and parameters used for minimap2 and other tools). Users have the option to use the LAST aligner [15], or can choose any aligner that generates outputs compliant with SAM [16] or MAF (http://genome.ucsc.edu/FAQ/FAQformat.html#format5) formats. Alignment data are filtered to keep the primary alignments of each read for further analysis. We describe the details of each analysis stages in the following sections. The simulation stage takes a reference transcriptome as input (FASTA/FASTQ formats). Reads will be simulated from the input transcriptome sequences. For each simulated read, it selects a transcript based on expression profiles calculated in the characterization stage. It then extracts a sequence from that transcript based on the length distribution model, and applies IR and error models to modify the extracted sequence. The simulation stage utilizes all information from the characterization step to produce *in silico* reads along with comprehensive information on introduced errors. The modularity of the Trans-NanoSim enables users to run it in their desired mode for detection of IR events or quantification of transcripts expression levels.

Trans-NanoSim is developed in Python and is freely accessible at https://github.com/bcgsc/NanoSim (Licence: GPL-3). It is platform independent and requires the following python packages to work: Six, numpy (Tested with version 1.10.1 or above), HTSeq, scipy (Tested with verson 1.0.0), scikit-learn (Tested with version 0.20.0). Reading and analyzing GTF/GFF3 files requires GenomeTools [17].

### Characterizing the length distribution

NanoSim [12] utilizes an empirical cumulative density function to simulate the length distribution of reads. In the current version of the pipeline, we improve on that by using kernel density estimation (KDE), which captures important patterns in the read length distributions, and avoids overfitting. We also remove the binning strategy in simulating the align ratio on each reads, resulting in a smoother simulated read length distribution. Theoretically, nanopore transcriptome sequencing can yield reads of the same length as the original mRNA molecule. However, in practice, ONT reads are generally shorter than the reference transcript they are derived from, which is attributable in part to RNA degradation and the tendency of reverse transcriptase molecules to disengage before reaching the 5’ end of the template RNA [18–19]. Therefore, it is crucial to consider the length of the reference transcript when simulating the length distribution of synthetic ONT reads. In order to achieve this, we utilize a two dimensional KDE model, and measure the length of an ONT read relative to the length of the reference transcript. Furthermore, based on alignment results, soft-clipped regions at the beginning and at the end of each read are also considered for length distribution analysis. We refer to these regions as head and tail in this study and follow the same approach to separately model the length distribution of these flanking regions.

We note that, the percentage of antisense sequences in cDNA and direct RNA sequences may be substantially different. To capture this information, Trans-NanoSim automatically calculates the percentage of reads that are on the positive strand with respect to the direction of transcription of the reference transcript, and applies this ratio in the simulation stage to maintain the strand ratio in simulated reads.

### Intron Retention module

The IR module of Trans-NanoSim detects and models these events using input ONT transcriptome reads. It uses alignments to intronic regions to calculate Markov chain transition probabilities between the states of spliced and retained introns, given the state of the previous intron. This feature is part of the characterization phase by default, and requires a reference genome and its annotation in GTF/GFF3 format. However, users may disable this option if they desire to do so. The module outputs comprehensive information on the location of the detected IR events based on input ONT reads. This functionality can be invoked in a standalone mode (*detect_ir*), enabling users to only detect and model IR events without characterizing or simulating reads.

### Transcript abundance quantification

We incorporate a pipeline (https://github.com/jts/nanopore-rna-analysis) to estimate transcript abundance based on reference transcriptome alignments (this functionality is provided, courtesy of Dr. Jared Simpson, personal communication). It relies on minimap2 [7] with -*p0* flag to retain all secondary mappings, and then utilizes an expectation-maximization approach similar to RSEM [20], Kallisto [21], and Salmon [22] to handle multi-mapping reads. It is a standalone module (*quantify*) which outputs transcript abundance in TPM values. The simulation stage of our pipeline relies on expression profiles to produce *in silico* reads.

### Characterizing the error modes

It was previously demonstrated that statistical modeling of error patterns in long nanopore reads can be useful in simulating reads mimicking the sequencing platform [12]. In Trans-NanoSim, we build on the same mixture models to deal with transcriptome reads as these patterns are shared between different library preparation methods and datasets. According to their alignment against the reference, reads are classified into two groups: aligned and unaligned. For each group, we consider specific characterization and modeling approaches. As for the aligned reads, we consider their aligned bases for further error rate analysis. The length of indels and mismatches are drawn from Weibull/Geometric and Poisson/Geometric mixture models, respectively. We also calculate the transitional probability between each two consecutive error types using a Markov chain model. We re-implemented the model fitting function of NanoSim in Python (formerly in R), and allowed multi-threading to expedite the fitting process. Unaligned reads may provide crucial information about the nature of ONT sequencing experiments, and thus we chose to model the length distribution of the unaligned reads as well. For this purpose, we extract sequences from reference transcripts based on their length distribution and apply an arbitrarily high error rate (90%).

### Datasets used for analysis

The datasets supporting the conclusions of this article are available from following sources. ONT cDNA and direct RNA sequencing reads of human NA12878 transcriptome are available under RNA project of Nanopore WGS Consortium (https://github.com/nanopore-wgs-consortium/NA12878/blob/master/RNA.md). ONT MinION RNA-seq of single mouse B1a cells are accessible by BioProject ID PRJNA339767 (https://www.ebi.ac.uk/ena/data/view/PRJNA339767) [4]. Please see **Supplementary Note 1** for details of datasets used in this study.

