## Supplementary information for "Trans-NanoSim characterizes and simulates nanopore RNA-seq data"

### Supplementary materials for
### "Trans-NanoSim characterizes and simulates nanopore RNA-seq data"

Saber Hafezqorani, Chen Yang, Ka Ming Nip, René L Warren, Inanc Birol

#### Supplementary Note 1: Datasets

We used publicly available direct RNA and cDNA ONT sequencing reads describing human and mouse transcriptomes provided by the Nanopore WGS Consortium RNA project and Byrne et al, respectively. From the Nanopore WGS Consortium – RNA project there are 30 runs (flowcells) of direct RNA reads sequenced on the Oxford Nanopore MinION device from 5 different centres. In addition, there are 12 cDNA read datasets sequenced using 1D ligation kit (SQK-LSK108) using the R9.4 chemistry. We also downloaded and used cDNA sequences from mouse transcriptome provided in the Byrne et al manuscript. The details of datasets we used for all analysis presented in our manuscript are as follows:

Short term in manuscript: Human cDNA dataset
- Centre: Birmingham (Bham) – Run 1
- Sample Type: 1D cDNA
- Kit: SQK-PCS108
- Pore Chemistry: R9.4
- Number of reads: 890503
- Reference: https://github.com/nanopore-wgs-consortium/NA12878/blob/master/RNA.md
- Human reference transcriptome used for analysis: Homo_sapiens.GRCh38.cdna.all.fa (Ensembl)

Short term in manuscript: Human direct RNA dataset
- Centre: Birmingham (Bham) – Run 2
- Sample Type: RNA
- Kit: SQK-RNA001
- Pore Chemistry: R9.4
- Number of reads: 221795
- Reference: https://github.com/nanopore-wgs-consortium/NA12878/blob/master/RNA.md
- Human reference transcriptome used for analysis: Homo_sapiens.GRCh38.cdna.all.fa (Ensembl)

Short term in manuscript: Mouse cDNA dataset
- Experiment: SRR5286960
- Sample Type: 1D cDNA
- Pore Chemistry: R9.4
- Number of reads: 104990
- Reference: https://www.nature.com/articles/ncomms16027
- Mouse reference transcriptome used for analysis: Mus_musculus.GRCm38.cdna.all.fa (Ensembl)

### Supplementary Note 2: Simulating reads from human and mouse reference transcriptomes

**Trans-NanoSim:** For each separate experiment, we used the empirical reads mentioned in the above dataset section to train Trans-NanoSim. We used mouse and human reference transcriptomes as input along with these reads, in the characterization phase. By default, Trans-NanoSim uses minimap2 (v2.17-r941) to align reads to reference transcriptome. Later, based on these alignments, Trans-NanoSim calculates the length distribution of reads as well as their error profiles. It further detects and models the intron retention events by default. We used the *quantify* module of the pipeline to quantify expression levels of the reads. Next, we used all these information in the simulation phase of the pipeline to generate the exact number of synthetic reads as raw reads. All the benchmark results presented in this manuscript are done using release v2.4-beta. Detailed information about set parameters and input files used in each step is as follows:

1. Running the characterization stage of Trans-NanoSim with the following input files:
    a. *–i*: Input ONT cDNA / directRNA reads in FASTQ/FASTA formats
    b. *–rg*: Reference genome in FASTA
    c. *–rt*: Reference transcriptome in FASTA
    d. *–annot*: Annotation file in ensembl GTF/GFF formats
    e. *–ga*: Genome alignment of input reads (note that it is optional. If you don't provide it, the pipeline will automatically align them to genome using minimap2 as well)
    f. *–ta*: Transcriptome alignment of input reads (note that it is optional. If you don't provide it, the pipeline will automatically align them to transcriptome using minimap2 as well)
2. Running the quantification module with the following input files:
    a. *–i*: Input reads for quantification (The same one we used in characterization stage)
    b. *-rt*: Reference transcriptome in FASTA
3. After getting both transcript expression levels and read profiles generated in the characterization stage, we run the simulation stage with the following input files:
    a. *–rt*: Reference transcriptome in FASTA
    b. *–rg*: Reference genome in FASTA
    c. *–e*: Expression profiles in *.tsv* format quantified in previous step
    d. *–c*: Location and prefix of read profiles generated in characterization step
    e. *–n*: Number of reads we want to simulate

For the full set of options available in each step and example runs, users may refer to the comprehensive readme file in GitHub repository: https://github.com/bcgsc/NanoSim

Options used for alignment tools:

- Alignment to reference transcriptome using minimap2: `minimap2 --cs -ax map-ont`
- Alignment to reference genome using minimap2: `minimap2 --cs --MD -ax splice`
- Alignment to reference transcriptome/genome using LASTAL:
    - Create indexes: `lastdb ref_genome` *`ref_g`*
    - Align: `lastal -a 1 -P` *`num_threads`*

**<u>DeepSimulator:</u>** We downloaded and used the latest released version of the DeepSimulator pipeline (v1.5) from the GitHub repository (https://github.com/lykaust15/DeepSimulator). In order to generate synthetic human cDNA/direct RNA and mouse cDNA reads, we used the reference transcriptome (human and mouse, respectively) as input to the tool for the simulation purpose. This new version of the DeepSimulator allows users to define the average length of the synthetic reads. To provide a fair comparison, we first calculated the average length of raw ONT transcriptome reads mentioned in the previous section and provided that average length in *-l* option of the DeepSimulator to maintain the mean length in simulated reads. We used Guppy CPU as the Basecaller. Other parameters such as the length distribution pattern were kept as default. We simulated the exact same number of synthetic reads as in raw experimental reads described in the dataset section. The following list describes options we used to simulate reads with DeepSimulator.

a. –*i*: We used reference transcriptome as an input for the Pipeline
b. –*o*: Defines the output directory of the simulated reads
c. –*B*: Defines the basecaller to be used. We ran the pipeline twice, one with Guppy CPU and one with Albacore.
d. –*l*: Defines the mean length of the synthetic reads. For this as described above, we first calculated the average length of empirical reads and used it here as input length
e. –*n*: Number of simulated reads we want to get

All simulation runs with Trans-NanoSim and DeepSimulator were done on cluster nodes running on Centos 6.7 system with 38 Intel Xeon E5-2650 CPUs and 380 GB of Memory.

### Supplementary Table 1

**Table S1: Error rates in empirical and simulated reads.** We aligned the simulated reads generated by Trans-NanoSim (TNS) and DeepSimulator (DS) back to the reference transcriptome and evaluated the rate of error events in simulated reads generated by Trans-NanoSim and DeepSimulator along with their raw experimental read counterparts (Raw). The results demonstrate that in all three datasets, reads generated by Trans-NanoSim contain similar error rates as empirical reads whereas error profiles of DeepSimulator deviate further away from experimental reads. Results indicate that deletion rates are usually higher in nanopore reads compared to mismatch and insertion events and we demonstrated that Trans-NanoSim is able to successfully capture this pattern. However, it seems like that DeepSimulator assumes all three error types to have similar rates and does not reflect the true patterns in empirical reads.

| % of the all bases having | Human cDNA | | | Human direct RNA | | | Mouse cDNA | | |
|---|---|---|---|---|---|---|---|---|---|
| | Raw | TNS | DS | Raw | TNS | DS | Raw | TNS | DS |
| Mismatch | 4.39 | 4.17 | 3.41 | 3.79 | 3.87 | 3.41 | 1.63 | 1.61 | 3.32 |
| Insertion | 4.42 | 4.24 | 2.58 | 3.84 | 3.28 | 2.58 | 1.62 | 1.53 | 2.62 |
| Deletion | 6.49 | 5.68 | 3.20 | 6.62 | 6.08 | 3.20 | 5.58 | 5.39 | 3.01 |
| Total | 15.30 | 14.10 | 9.19 | 14.26 | 13.34 | 9.20 | 8.84 | 8.53 | 8.96 |

### Supplemental Figures

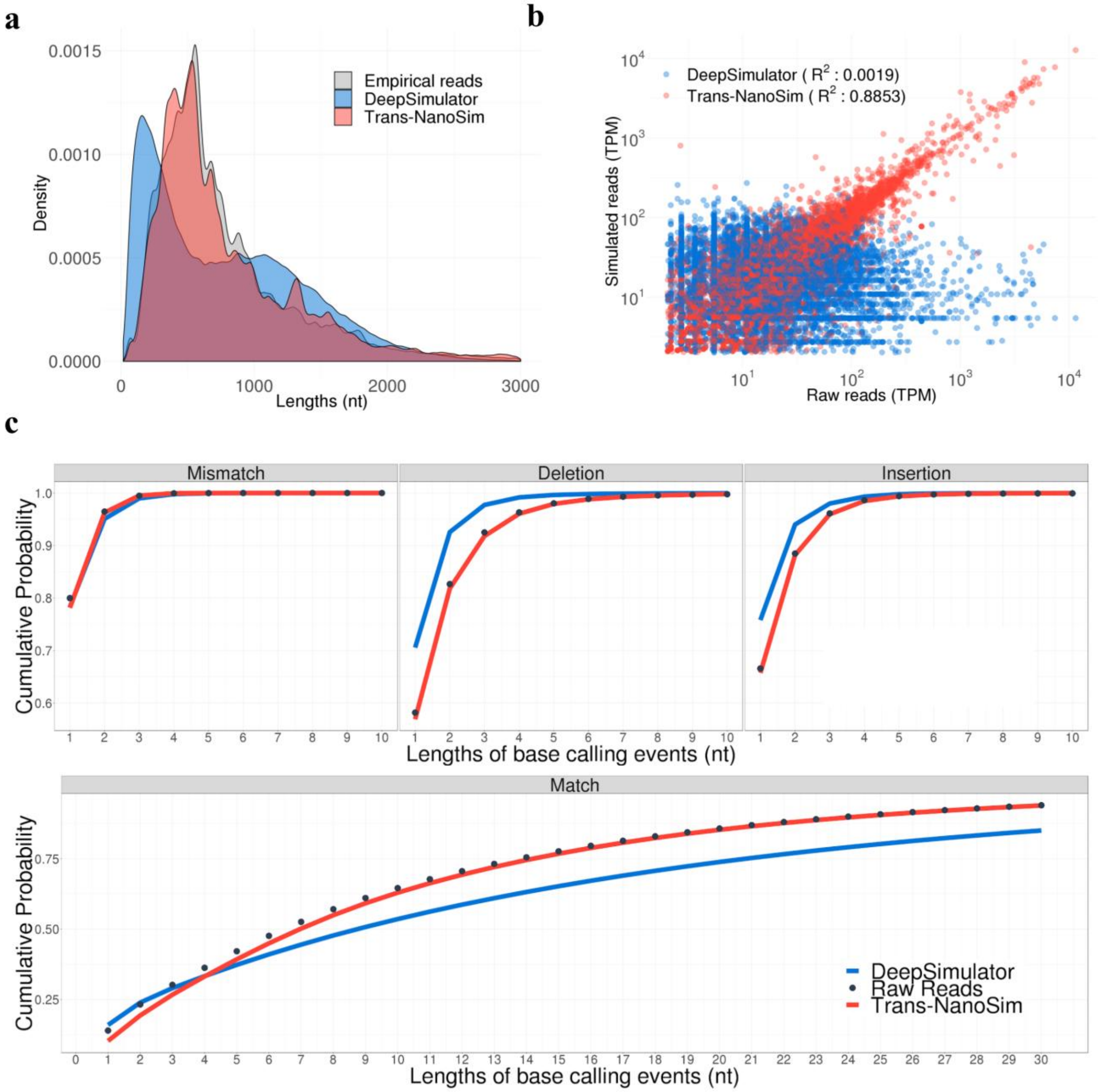


**Figure S1: Benchmarking Trans-NanoSim and DeepSimulator on the human direct RNA dataset. a, a,** Comparison of length distributions of experimental reads and simulated reads generated by Trans-NanoSim and DeepSimulator. **b,** Transcript expression levels measured from simulated reads versus the same measured from experimental reads. **c,** The length of consecutive match/error bases of empirical and simulated reads, as indicated.

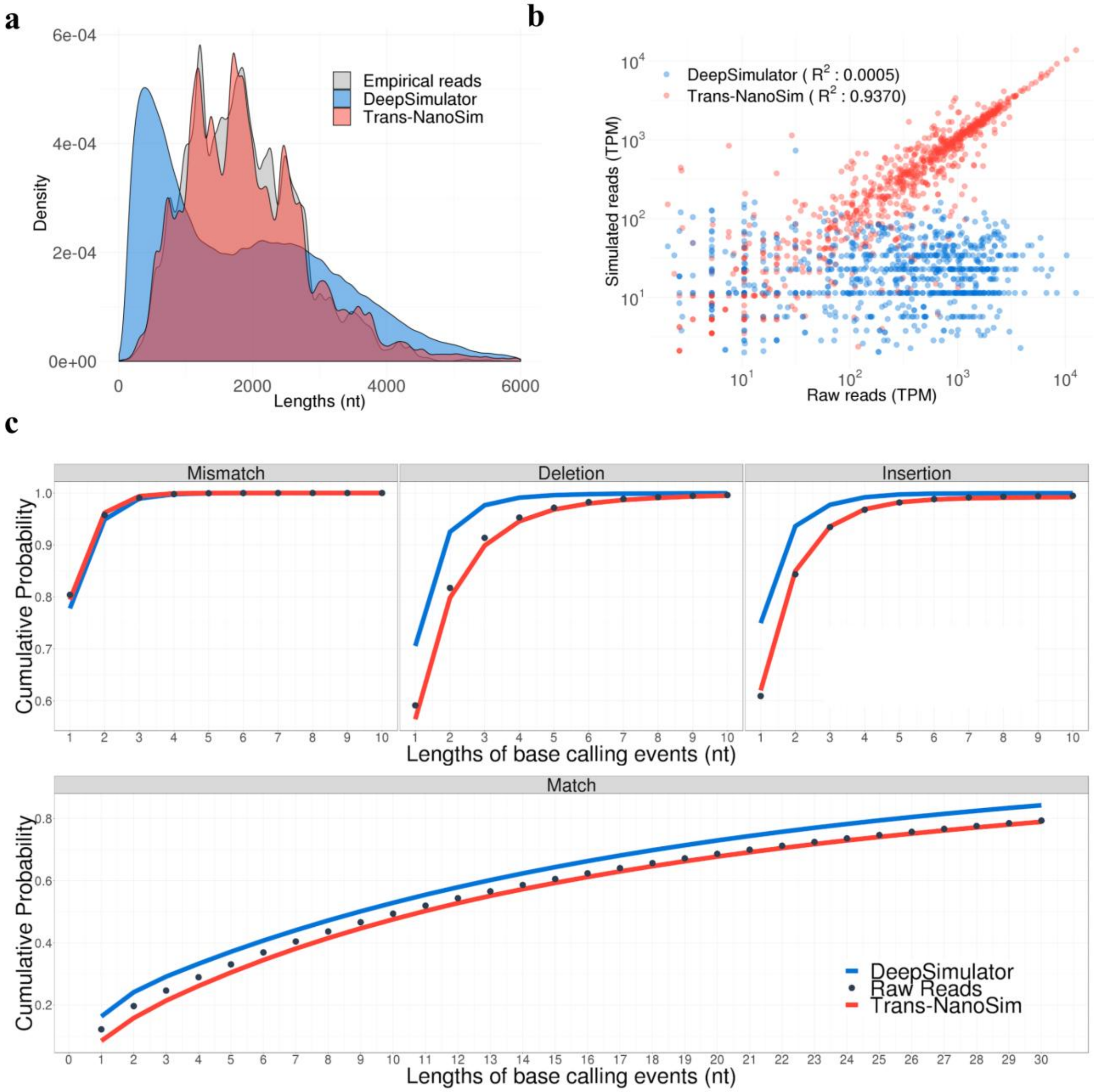


**Figure S2: Benchmarking Trans-NanoSim and DeepSimulator on the mouse cDNA dataset. a, a,** Comparison of length distributions of experimental reads and simulated reads generated by Trans-NanoSim and DeepSimulator. **b,** Transcript expression levels measured from simulated reads versus the same measured from experimental reads. **c,** The length of consecutive match/error bases of empirical and simulated reads, as indicated.

#### Supplementary Note 3: Statistical test of read length distributions

We statistically tested the length distribution of synthetic reads generated by Trans-NanoSim through ordinary nonparametric bootstrapping 1000 times (`boot` command in R) and calculating the 95% confidence interval. Figure S 3-5 illustrates the bootstrapping results on human cDNA, human direct RNA and mouse cDNA datasets, respectively.

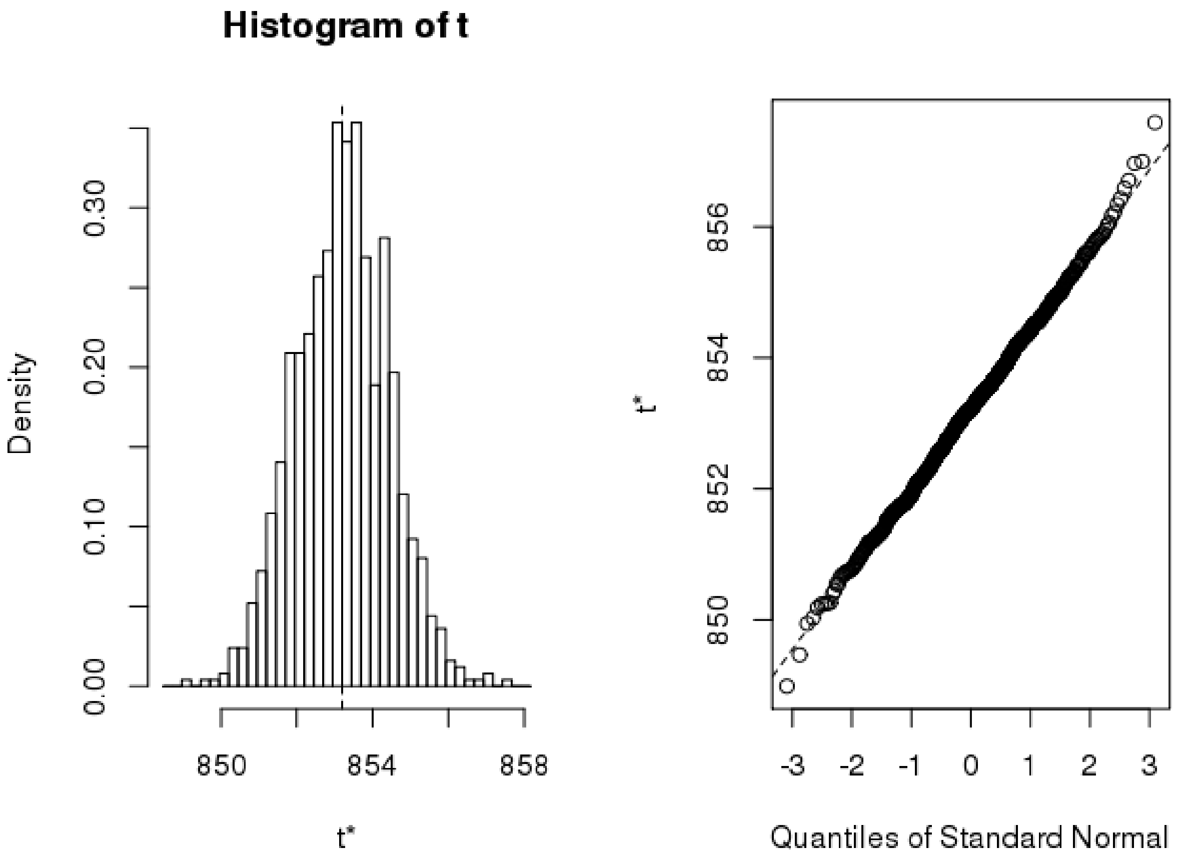


**Figure S3: Bootstrapping results for the human cDNA dataset.** The average length of the original read set is 853 (SD=1) nt. The 95% confidence interval is between 850 and 855 nt. The x-axis in the left plots (t*) is the average length of reads in each bootstrapping sample set.

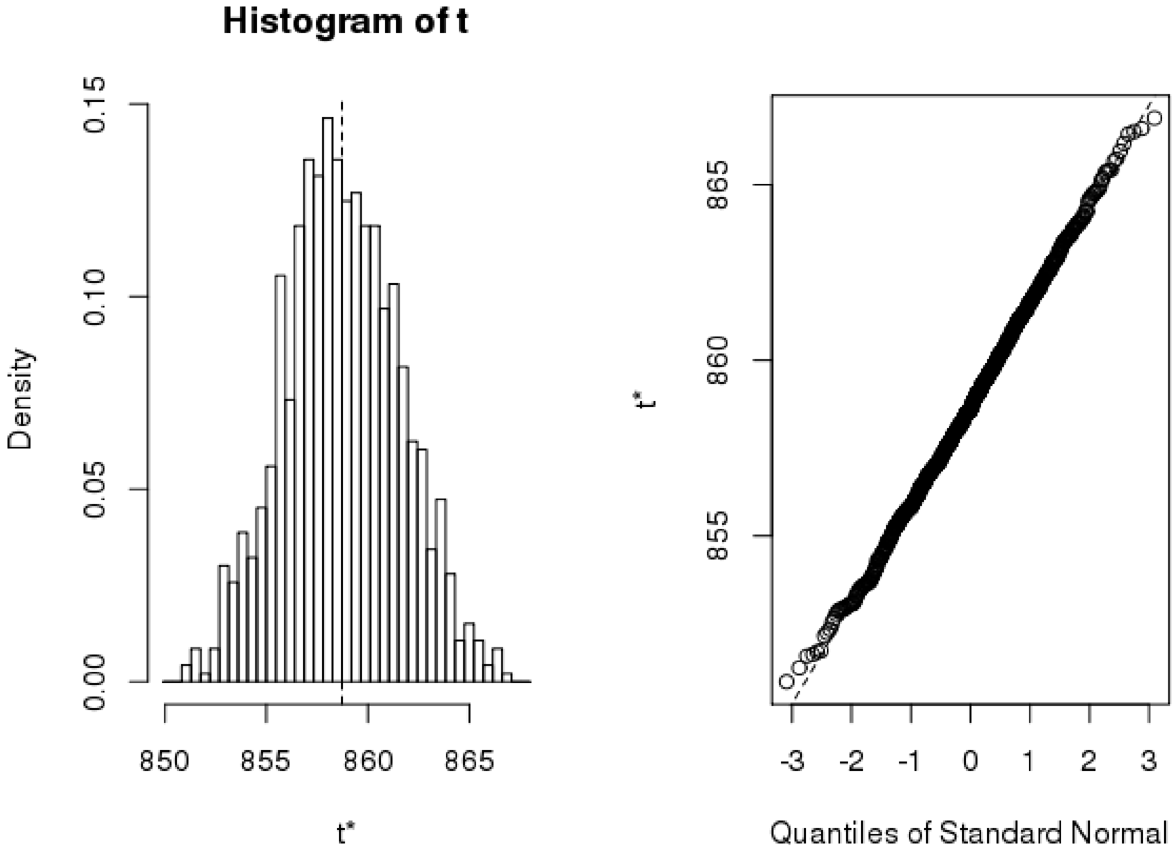


**Figure S4: Bootstrapping results for the human direct RNA dataset**. The average length of the original read set is 858 (SD=3) nt. The 95% confidence interval is between 853 and 864. The x-axis in the left plots (t*) is the average length of reads in each bootstrapping sample set.

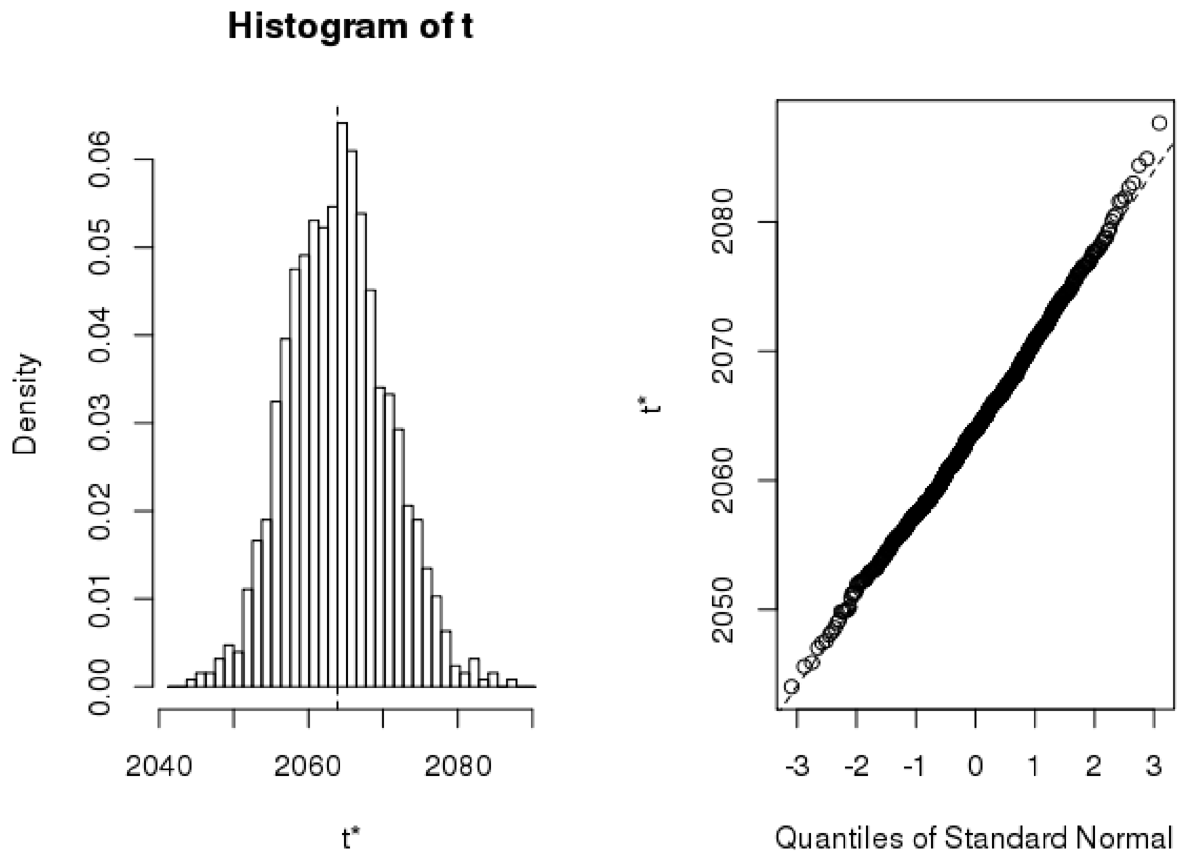


**Figure S5: Bootstrapping results for the mouse cDNA dataset**. The average length of the original read set is 2063 (SD=7) nt. The 95% confidence interval is between 2051 and 2077. The x-axis in the left plots (t*) is the average length of reads in each bootstrapping sample set.
